# Sperm Toolbox – A selection of small molecules to study human spermatozoa

**DOI:** 10.1101/2023.05.02.539126

**Authors:** Franz S. Gruber, Anthony Richardson, Zoe C. Johnston, Rachel Myles, Neil Norcross, David Day, Irene Georgiou, Laura Sesma Sanz, Caroline Wilson, Kevin D. Read, Sarah Martins da Silva, Christopher L. R. Barratt, Ian H. Gilbert, Jason R. Swedlow

## Abstract

Male contraceptive options and infertility treatments are limited. To help overcome these limitations we established a small molecule library as a toolbox for assay development and screening campaigns using human spermatozoa. We have profiled all compounds in our automated high-throughput screening platform and provide the dataset, which allows direct side-by-side comparisons of compound effects and assays using live human spermatozoa.

## Main

One of the grand challenges for science and society is the development of novel contraceptives that are safe and deliver control of reproduction to men and women all over the world. In parallel, a deeper understanding of sperm maturation, motility, fertilization, and possible interventions for male infertility is required to counter declining birth rates. Recently, the application of advanced technologies has helped move the human fertility field forward, revealing new potential targets for both contraception and fertility, and new compounds as contraceptive candidates (reviewed by ^1-3^). A key part of this effort is the development of mechanistic insight and drug candidates that affect the behaviour and properties of human spermatozoa ^4-7^. Towards this end, we have recently developed an automated live human sperm phenotypic, target-agnostic screening platform that can profile 1000s of compounds and identify modifiers of sperm function ^8,9^. These effects can be either positive or negative and therefore are starting points for either fertility treatments (including medical assisted reproductive technologies) or contraceptive development. An advantage of using a high-throughput platform is the ability to efficiently compare many compounds under the same assay conditions, within a short amount of time.

Target-agnostic phenotypic screening can be enhanced by assaying libraries of compounds with known targets. The resulting reference datasets can suggest mechanism(s) of action and target identification for candidate compounds during a screening campaign. To facilitate our own, and others, drug discovery efforts in contraception and infertility, we assembled a small library containing 84 small molecules, which we have coined the Sperm Toolbox (STB). The idea of the STB is in concordance with similar efforts e.g. the MMV Pandemic Response Box ^10^, the SGC Epigenetic Chemical Probe Collection ^11^, or the Drug Repurposing Hub ^12^ that facilitate deeper understanding of biology or pathology or repurpose approved drugs. We tested the STB compounds in multiple functional sperm assays, as well as more general cytotoxicity and aqueous solubility assays (Figure 1A). The STB has been assembled to contain previously published compounds and compounds detected in our own screening campaigns ^8,9^. In addition, we included a few compounds which have been inferred from patent applications or studies of proteins potentially relevant for sperm function (Figure 1B). In constructing the STB, we have focussed on compounds reported in the literature that affect human systems, but compounds tested in relevant animal models, cell-based, or biochemical systems have also been included. Published analyses of the compounds selected in the STB have indicated effects on crucial sperm functions e.g. motility, capacitation, acrosome reaction, Ca^2+^ signalling, viability, or related to other processes (e.g. spermatogenesis, liquefaction, or ejaculation) (Figure 1C). Compound annotations are included in the STB dataset and indicate that Enzymes/Kinases, Receptors, channels and transporters are the most represented compound classes. Using information from the Drug Repurposing Hub ^12^, we generated a protein-protein interaction network of targets related to STB compounds (Figure 1E). This network consists of multiple subclusters related to functions in spermatozoa (Figure 1F) and other biological processes, functions, and pathways (Figure 1G).

**Figure 1.**
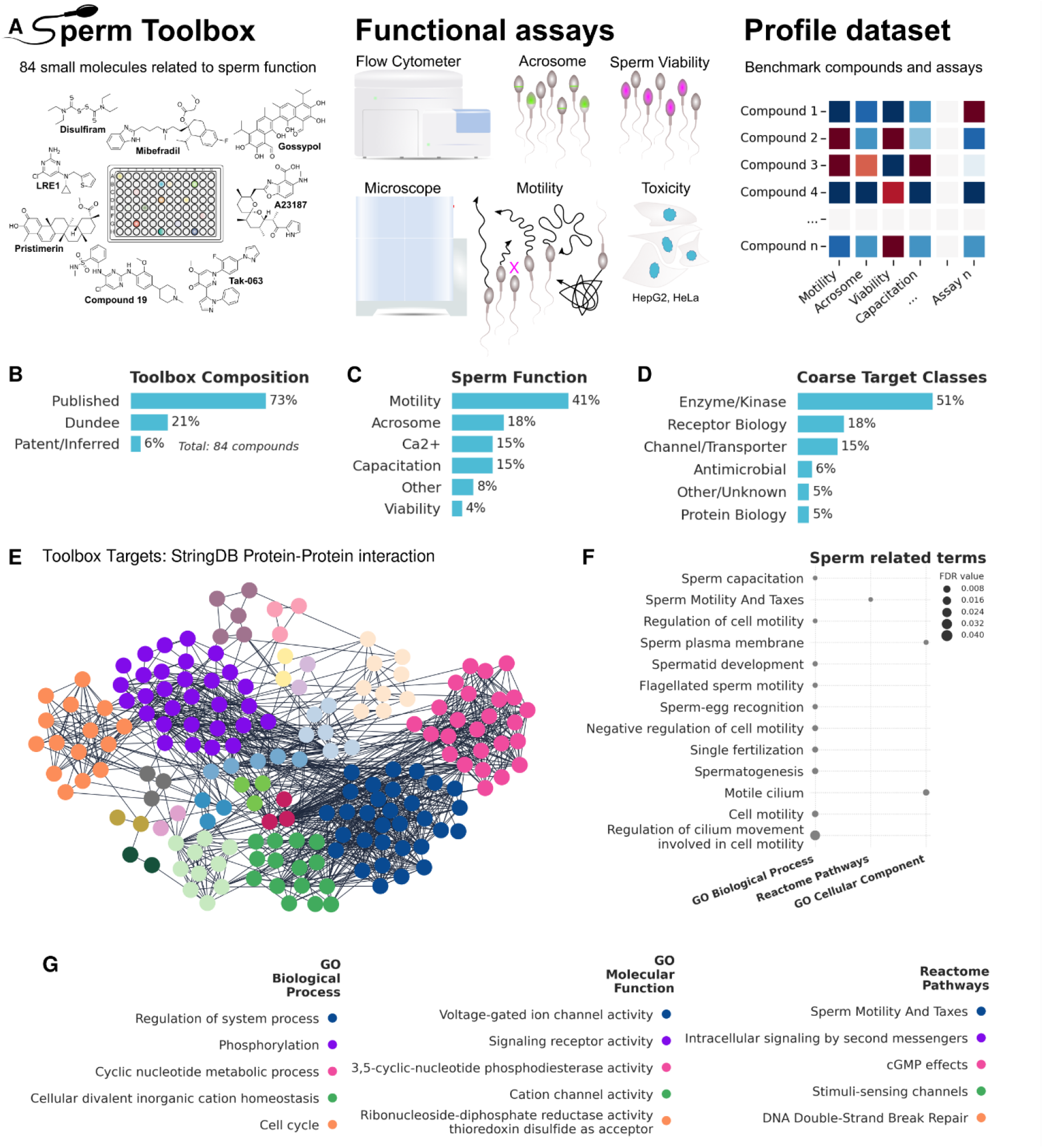
(A) The Sperm Toolbox is a collection of 84 small molecules with published, observed or inferred effect on multiple biological processes related to the biogenesis or function of spermatozoa. Human sperm cells have been tested in various assays, which allows comparison of compounds under the same experimental conditions. (B) Summary of the sperm toolbox compound compositions. (C) Summary of annotated (i.e. published, observed, inferred) sperm function of Toolbox compounds. (D) Coarse classification of Sperm Toolbox target classes. (E) StringDB Protein-Protein interaction networks (Full network, confidence 0.7) of putative targets of Sperm toolbox compounds. Protein targets have been inferred from the repurposing hub. Network has been clustered using MCL clustering in Cytoscape. (F) Sperm related terms (GeneOntology, Reactome) within the network shown in (E). Marker size indicates false discovery rate (FDR). (G) Enriched terms within the biggest subclusters in (E). Redundant terms have been filtered using Cytoscape (cutoff 0.5). Color matches subcluster color in (E).

To compare the effects of STB compounds on specific sperm functions, we screened them in multiple phenotypic functional assays. Each sperm function assay has been repeated in multiple donor pools (see Material and Methods), where each pool consists of cells donated from at least 3 different donors to address donor to donor variability. Each compound has been tested at two concentrations, 10 and 30 μM, and normalized to vehicle controls (DMSO). We have modified our assays from the previously published workflow ^8^ by increasing incubation time and introducing conditions that support capacitation, which is a sperm maturation process required for fertilization. These changes expand the range of targets that we can explore in our assays. Our live-sperm-based high-throughput assays use an automated microscope to record short timelapse movies and can run in different modes. The first mode, denoted ‘motility assay’, utilises non-capacitating assay conditions that model the state of spermatozoa after ejaculation. The second mode, denoted ‘capacitation assay’, utilises conditions which support capacitation (Extended Figure 1A). Both modes allow sperm motility to be measured. In addition, the capacitation assay also scores hyperactivation (i.e. distinct movement patterns concomitant with capacitation which are crucial for fertilization)

The acrosome assays utilize a high-throughput flow cytometer to score for compounds inducing the acrosome reaction, using peanut agglutinin conjugated to a fluorescent dye to label the inner acrosomal membrane. As with the motility assays, we have developed two different screening modes, named ‘acrosome 1’ and ‘acrosome 2’ (Extended Figure 1A). Both modes are capable of measuring induction of the acrosome reaction. However, in the acrosome 2 assay, a mixture of physiological agonists of the acrosome reaction is used ^13^. This addition allowed measurement of inhibition of acrosome reaction.

We also tested the effect of STB compounds on sperm viability using a complementary live-cell flow-cytometer approach and propidium iodide, which does not permeate intact cell membranes (Extended Figure 1B).

Finally, we also profiled each STB compound for general cytotoxicity in two cell-lines (HeLa and HepG2). The HeLa-based assay uses the cell painting assay ^14^, while the HepG2 assay utilizes a resazurin based viability read-out ^15^. To ensure usability of STB compounds, we also measured aqueous solubility. An overview of additional assays is given in Extended Figure 1C.

This multi-assay compound profiling dataset reveals a broader picture of the action of individual compounds and helps find phenotypic similarities induced by compounds. Using unsupervised clustering we observed clusters of compounds with a significant reduction of sperm motility and/or increased amounts of acrosome reacted cells (e.g. Alexidine dihydrochloride, Tafenoquin or A23187) (Figure 2A). Some of these compounds also show significant levels of cytotoxicity in HeLa and/or HepG2 cells as well as decreased sperm viability suggesting that their effect on motility and acrosome reaction is due to toxic effects on spermatozoa. However, other clusters show a significantly increased acrosome reaction and hyperactive motility but do not show effects on sperm viability or general cytotoxicity (e.g. LRRK2-IN-1, Clofarabine, Linsitinib, Trequinsin). One example, LRRK2-IN-1, which enhances motility (progressive and hyperactivated), has been described as a selective inhibitor of LRRK2 kinase, which is involved in Parkinson’s disease ^16^. This compound shares some structural similarities with two kinase inhibitors in the STB, which decrease sperm motility: Compound 19, described as a potent Testis-Specific Serine/Threonine Kinase 2 (TSSK2) inhibitor ^17^ and CZC 25146, annotated as a potent LRRK2 inhibitor.

**Figure 2.**
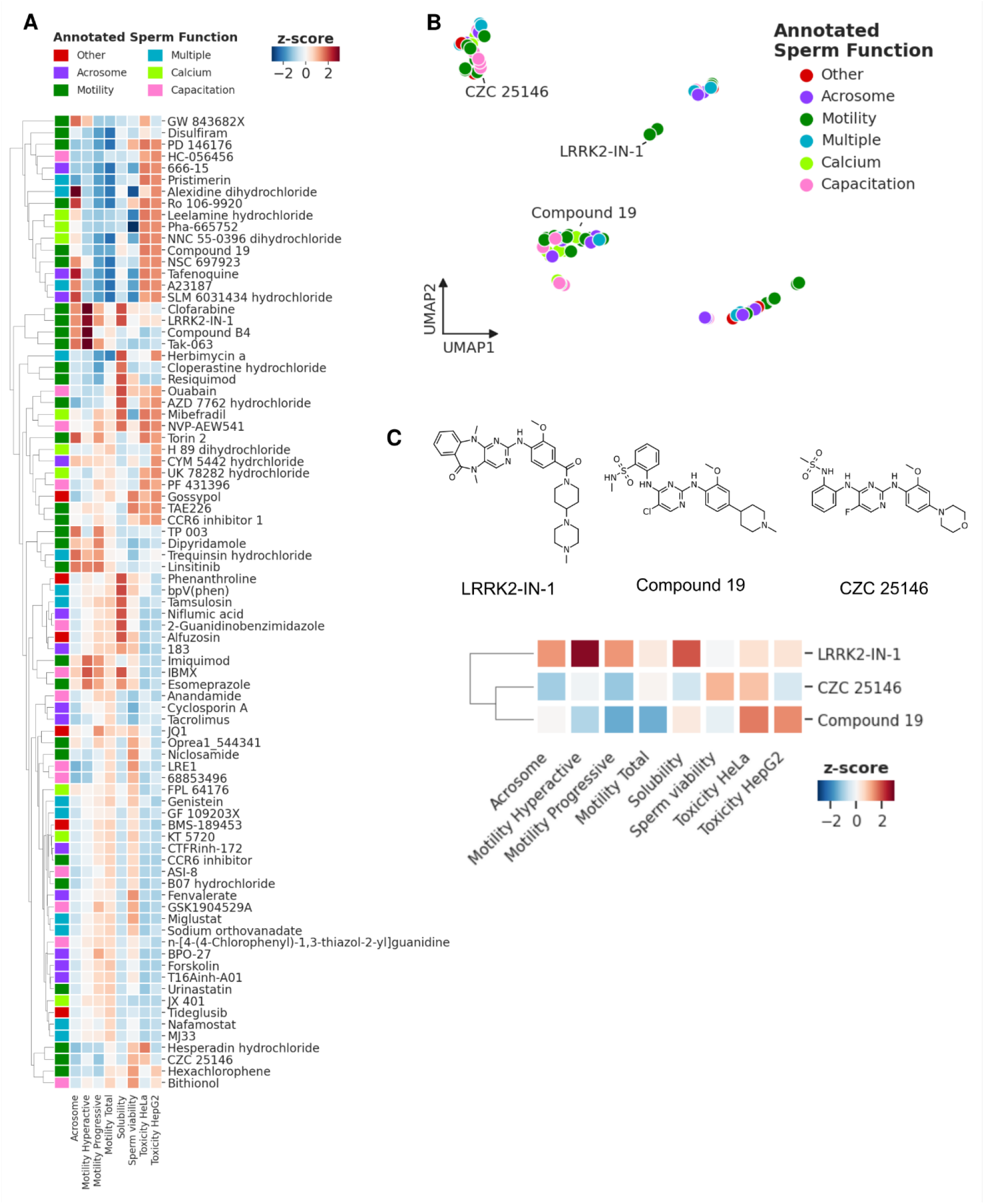
(A)Heatmap showing standardized results (z-score) for every Toolbox compound screened at 30 μM in selected assays: Acrosome 1 (3h with compound then induction with acrosome reaction using an agonist mix), Motility Hyperactive (30 min capacitation then 30 min incubation with compound), Motility Progressive (3h incubation with compounds under non-capacitating conditions), Motility Total (3h incubation with compound under non-capacitating conditions)), Solubility (aqueous solubility of compounds after 24h incubation), Sperm viability (3h with compound then propidium iodide assay), Toxicity HeLa (24h compound incubation then cell painting assay), Toxicity HepG2 (70h incubation with compound then resaruzin based assay). Heatmap color indicates decrease (blue) or increase (red). (B)UMAP plot showing groups of compounds with similar assay profile. Color indicates annotated sperm function. (C)Structures and zoom-in of three examples related to LRRK2 kinase.

Interestingly, we observed that a number of compounds show weak or no effects in our assays (Extended figure 2; grey datapoints). Given these compounds were sourced from publications, patents, and screening systems that report activity in sperm, this highlights the need for standardisation of conditions in phenotypic assays based on human sperm. The STB, and the use of high-throughput assays allows the profiling of compounds using the same assay conditions (i.e. buffer system, incubation time and screening concentration) on a large number of human spermatozoa, thus providing reproducible, quantitative assay results. This combination also facilitates the comparison of the effects of the STB compounds between different physiological states (Extended Figure 1A&B), in a standardised, rapid, and quantitative manner that has not been previously possible in the field of sperm biology. This standardized approach provides an excellent starting point to establish control compounds and assay conditions for novel drug discovery campaigns related to contraception and infertility. We provide the complete dataset (see Supplement Information) as well as web interface for data analysis (https://spermtoolbox.shinyapps.io/spermtoolbox/).

This work represents the first version of the STB. We aim to release annual updates with newly published compounds modulating functions in spermatozoa. In our next release we aim to include compounds that were published during the collection of the data for this first release e.g., a sAC inhibitor TDI-11861 ^4^, a CatSper inhibitor RU1968 ^18^, a SLO3 inhibitor ^19^ or other contributions from the research community. The STB is a reference for compounds that modulate sperm function and can become an important resource for benchmarking new assays and new compounds. We aim to use the STB as a reference in our screens and to define as many factors targeted by STB compounds as possible that are critical for sperm function.

**Extended Figure 1.**
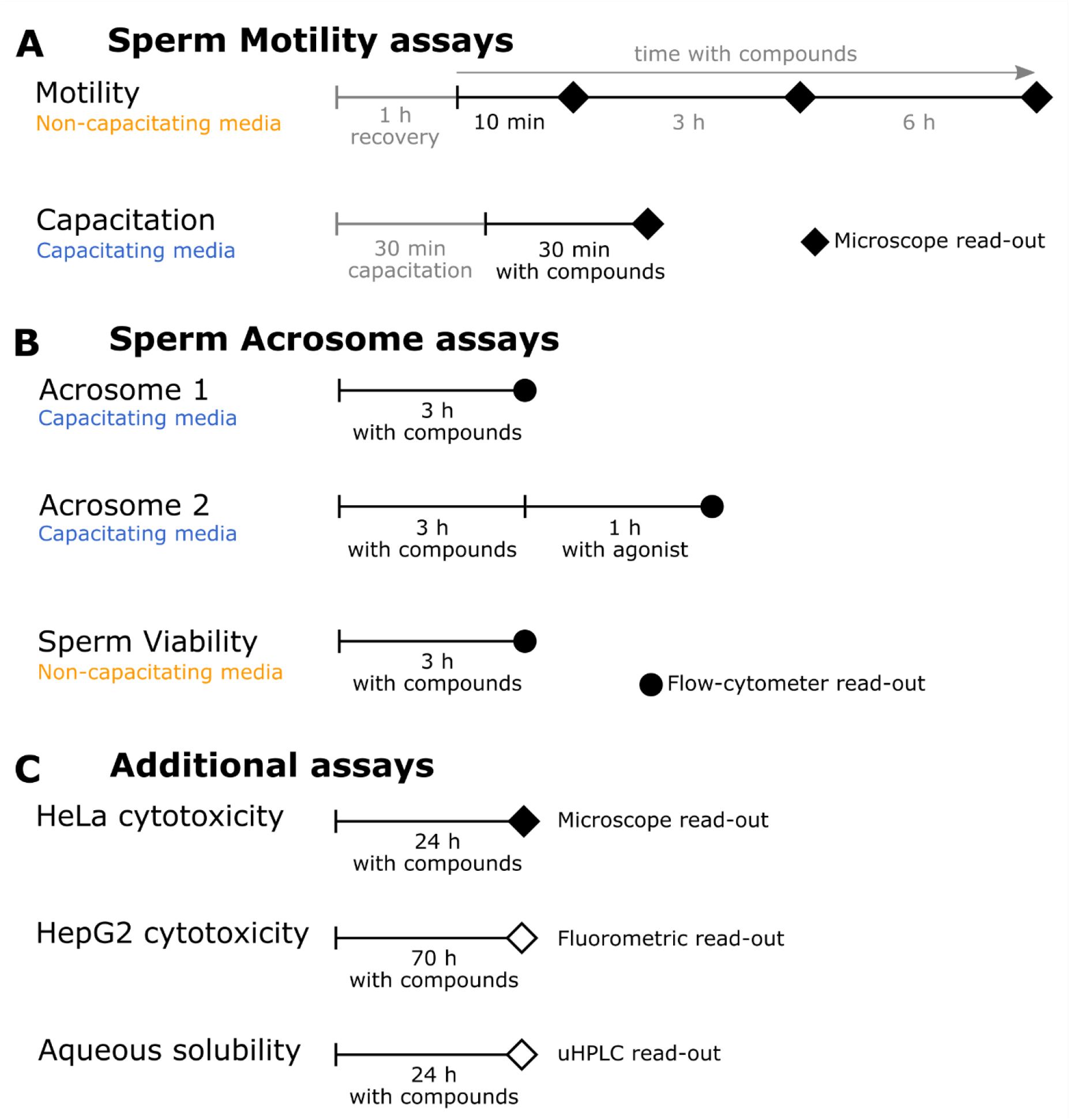
(A)Diagram showing sperm motility assays run on high-content microscope with compound incubation time/read-out time. Motility assays were run under non-capacitating conditions, with a 1 hour recovery phase after preparation of spermatozoa. Capacitation assays were run under capacitating conditions. Spermatozoa are capacitated for 30 min prior to compound incubation. (B)Diagram showing assays run on high-throughput flow cytometer to measure acrosome status and sperm viability. Incubation time with compound, agonist and read-out times are indicated. Acrosome assays run under capaciting conditions. Sperm viability assay run under non-capacitating condition. (C)Additional assays performed on sperm toolbox compounds.

**Extended Figure 2.**
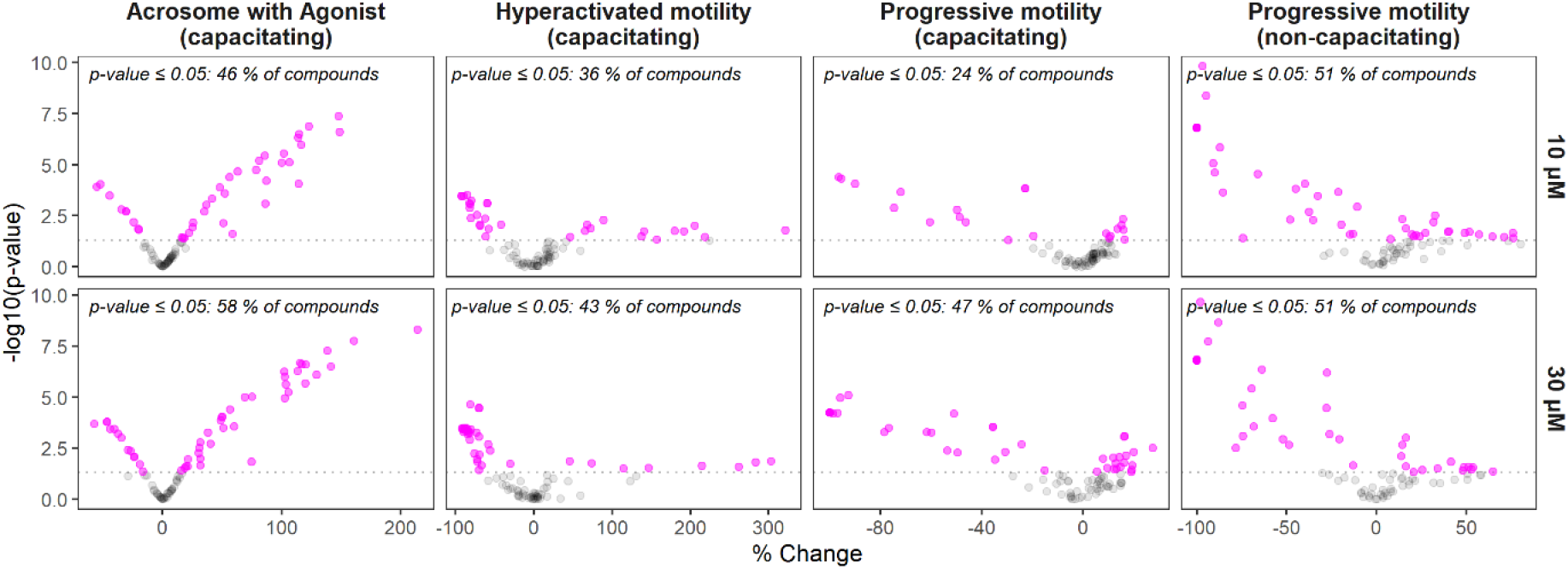
(A)Volcano plots of Acrosome, Capacitation and Motility assays. Upper panel indicates 10 μM, lower plot panels indicate 30 μM. Incubation times for assays are 3h with compound for Acrosome assay and Motility (non-capacitating) (PM = progressive motility), and 30 min for the Capacitation assay (HA = hyperactive motility, PM = progressive motility). Dotted line indicates significance levels of 0.05. Any point above dotted line is indicated in magenta. Buffer conditions are indicated below Plot title (capacitating vs. non-capacitating)

## Material and Methods

### Ethical approval

Written consent was obtained from each donor in accordance with under local ethical approval (University of Dundee, SMED REC 20/45).

### Sperm handling

Donated human spermatozoa with normal semen characteristics (concentration, motility; WHO 2010) from healthy volunteers with no known fertility problems were used in this study. Samples were obtained by masturbation after ≥ 48 h of sexual abstinence, liquefied for 30 min at 37 °C, and processed using density gradient centrifugation, to separate cells using 40/80% Percoll (Sigma Aldrich, UK) fractions. Every experiment has been performed on pooled spermatozoa from ≥ 3 donors each.

For experiments in non-capacitating conditions, we used a buffer system slightly modified from (Tardif et al., 2014) using Minimal Essential Medium Eagle (Sigma), supplemented with HEPES (1 M solution, Gibco), sodium lactate (DL-solution, Sigma), sodium pyruvate (100 mM solution, Gibco) and bovine serum albumin (7.5 % solution, Sigma).

For experiments in capacitating conditions, we used a buffer system slightly modified from the HTF from to a final composition of 97.8 mM NaCl, 4.69 mM KCl, 0.2 mM MgSO_4_, 0.37 mM KH_2_PO_4_, 2.04 mM CaCl_2_, 0.33 mM Na-pyruvate, 12.4 mM Na-lactate, 2.78 mM glucose, 21 mM HEPES, 25 mM NaHCO_3_ and 3 mg/mL BSA. All buffer components were supplied by Sigma-Aldrich unless otherwise stated and all buffer systems were adjusted to pH 7.4.

### Compound handling

Compounds were purchased from vendors indicated in supplementary file x. 10 mM stock solutions in DMSO were generated. Quality control of each compound has been performed (HPLC-MS and NMR).

Compound plates were generated using 10 mM stock solutions of each STB compound in Echo 384-LDV plates (LP-0200). Prior to running experiments, assay ready plates have been generated using Echo acoustic dispensers (550/555). Each compound was screened at 10 and 30 μM final concentration. Plate types used for assays: Motility, HeLa cell painting and Sperm viability (PerkinElmer 384-well PhenoPlate), Acrosome (Greiner 384-PP V-bottom plates).

### Motility assays

#### Motility assay (Non-capacitating)

Cells were rested for 1 h post density gradient centrifugation. 10 μL assay buffer was dispensed into assay ready plates, plates were put onto a shaker instrument for 30 seconds using 1,000 rpm, followed by a short pulse centrifugation. Plates were incubated for 15-30 mins at 37 °C prior to addition of 10 μL of pooled cells (at 1 M/mL concentration). Plates were then incubated in a Yokogawa CV7000 microscope set at 37 °C for 10 min prior to imaging. Imaging a 384-well plate requires ≤ 20 min using the following settings: 2 positions per well, 0.5 sec timelapse movies (3x binning; 11 ms exposure time, 22 ms interval time, 500 intervals). A second and a third timepoint was recorded after 3 h, and 6 h incubation at 37 °C. Data has been processed as described in Gruber et al., 2020.

#### Capacitation assay

Cells were capacitated for 30 min at 37 °C and 5 % CO2. 10 μL assay buffer was dispensed into assay ready plates, plates were put onto a shaker instrument for 30 seconds using 1,000 rpm, followed by a short pulse centrifugation. Plates were incubated for 15-30 mins at 37 °C and 5% CO2 prior to addition of 10 μL of pooled cells (at 1 M/mL concentration). This was followed by a 20 min compound incubation step at 37 °C and 5 % CO2. Plates were then incubated in our CV7000 microscope set at 37 °C and 5% CO2 for 10 min prior to imaging. Imaging a 384-well plate requires ≤ 20 min using the following settings: 2 positions per well, 0.5 sec timelapse movies (3x binning; 11 ms exposure time, 22 ms interval time, 500 intervals). Data has been processed as described in Gruber et al., 2020.

### Acrosome assays

In both experimental modes, 20 μL of pooled cells (at 1 M/mL concentration) were capacitated for 3 h in presence of compounds at 37 °C and 5% CO2. In mode a, cells were then stained with PNA-Alexa647 (ThermFisher, 1 mg/mL, final dilution 1:1000) and Hoechst dye for 30 min at 37 °C and 5% CO2, followed by 10 min fixation at RT by adding 20 μL of 4% paraformaldehyde (Sigma) in PBS. In mode b, a cocktail of progesterone, prostaglandin A and NH4Cl was added to all compound wells and incubated for an additional 1 h at 37 °C and 5% CO2 to induce acrosome reaction. This was followed by fixation and staining protocol as for mode a. In both modes fixatives and stains were removed and cells were resuspended in PBS, using a 405 platewasher. Plates were then measured on a Novocyte Advanteon high-throughput flow cytometer. Gating of data has been performed using the flow cytometer software to export counts of gated populations.

### Sperm viability assay

Cells were incubated with compounds for a total of 3h at 37 °C. 20 min prior, propidium iodide was added (1.5 mM solution, Sigma, final dilution 1:2000). Plates were analysed using a Satorius iQue Screener with settings to allow for a sampling time of ≤30 min per 384-well plate. Gating of data has been performed using the flow cytometer software to export counts of gated populations.

### HeLa cell painting

Cell were incubated with compounds for 24h. Cell painting was performed after Bray et al., 2016 with modifications suggested here (Cimini et al., bioRxiv 2022.07.13.499171; doi: https://doi.org/10.1101/2022.07.13.499171):

- 10 μL of MitoTracker (Life Technologies, M22426, 1 mM stock in DMSO as recommended by the vendor) is added to 50 μL of media (1:2000 final dilution)
- Phalloidin-Alexa594 (Life Technologies, A12381, 66 μM stock in methanol as recommended by the vendor, 1:500 final dilution)
- WGA-Alexa594 (Life Technologies, W11262, 5 mg/mL stock, 1:250 final dilution) was used
- DAPI (5 mg/mL stock; 1:1000 final dilution) was used to stain DNA
- 20 μL of staining solution was used
- PBS was used for washing steps

### HepG2 cytotoxicity assay

HepG2 assay was performed as described previously ^15^.

### Aqueous solubility

The aqueous solubility of test compounds was measured using UHPLC, as previously described in ^20^.

### Data analysis

For cell painting imaging data, plates of images were imported into OMERO ^21^. Cell segmentation and feature extraction were performed using CellProfiler 3 ^22^ and using instructions from ^14^ to normalize data. Timelapse data of motility assays was processed as previously described ^8^. Acrosome flow cytometry data was processed using NovoCyte Express (PerkinElmer), to gate out background debris, select for single cells and calculate percentage values of acrosome positive events. Sperm viability flow cytometry data was processed using iQue Forecyt (Satorius) to gate out background debris, select for single cells and calculate percentage values of propidium iodide positive events. For all sperm functional assays, data was normalized to DMSO control wells. Each plate had 16 DMSO wells. A median value of all 16 wells was used to calculated percentage change (pct_ch = ((value / DMSO_median) -1) * 100). For sperm viability we calculated fold-change (fc = 1 -(value / DMSO_median)). For each compound we calculated a p-value using a Welch’s t-test implemented in scipy (scipy.stats.ttest_ind) comparing a compounds effect to DMSO controls.

The following packages have been used for data analysis and visualization: R: Tidyverse, plotly, pHeatmap

Python: Pandas, NumPy, SciPy, Seaborn, Matplotlib, UMAP

## Acknowledgements

The authors acknowledge support by the Bill & Melinda Gates Foundation (INV-007117).

